# Multiple two-photon targeted whole-cell patch-clamp recordings from monosynaptically connected neurons in vivo

**DOI:** 10.1101/516708

**Authors:** Jean-Sébastien Jouhanneau, James F.A. Poulet

## Abstract

Although we know a great deal about monosynaptic connectivity, transmission and integration in the mammalian nervous system from *in vitro* studies, very little is known *in vivo*. This is partly because it is technically difficult to evoke action potentials and simultaneously record small amplitude subthreshold responses in closely (< 150 µm) located pairs of neurons. To address this, we have developed *in vivo* two-photon targeted multiple (2 – 4) whole-cell patch clamp recordings of nearby neurons in superficial cortical layers 1 to 3. Here we describe a step-by-step guide to this approach in the anesthetised mouse primary somatosensory cortex, including: the design of the setup, surgery, preparation of pipettes, targeting and acquisition of multiple whole-cell recordings, as well as in vivo and post-hoc histology. The procedure takes ∼ 4 hours from start of surgery to end of recording and allows examinations both into the electrophysiological features of unitary excitatory and inhibitory monosynaptic inputs during different brain states as well as the synaptic mechanisms of correlated neuronal activity.

## INTRODUCTION

Monosynaptic transmission underpins action potential generation and the flow of information within neural circuits. Over the last decades, *in vitro* approaches have provided an enormous amount of data on the connectivity rates and the electrophysiological and anatomical properties of synapses. More recently, a hybrid approach has been developed to link the neuronal function, measured in vivo, with the connectivity, measured in vitro (Cossell et al., 2015; Ko et al., 2011; Weiler et al., 2018). There is still, however, a large gap in our knowledge about monosynaptic transmission and membrane potential (V_m_) recordings of connected neurons in vivo.

In vivo approaches to identify connected pairs of neurons in the mammalian nervous system have typically performed electrophysiological recordings of multiple single neurons and examined the average response of one neuron to spontaneously occurring action potentials in another neuron. “Spike triggered averaging” of cortical neurons has been performed both with multiple extracellular recordings (Barthó et al., 2004; Csicsvari et al., 1998; English et al., 2017; Fujisawa et al., 2008; Reid and Alonso, 1995; Swadlow and Gusev, 2002) and a combination of extracellular and intracellular V_m_ recordings (Bruno and Sakmann, 2006; Crochet et al., 2005; London et al., 2010; Matsumura et al., 1996; Yu and Ferster, 2013). However, because it is not yet possible to record the activity of all neurons presynaptic to the cells of interest and cortical neurons can fire simultaneously, it is difficult to confirm whether correlated activity is the result of a direct synaptic connection between the two recorded neurons or input from a third, unrecorded neuron with similar firing dynamics. One approach to get around this problem is to have experimental control of action potential timing using single cell stimulation while simultaneously recording the evoked membrane potential (V_m_) response from a second neuron. While care has to be taken in concluding that any synaptic response is the result of a monosynaptic rather than polysynaptic input (Berry and Pentreath, 1976; Parker, 2010), this approach has been used in vivo to characterize the wiring and functional properties of synaptic connections in a number of non-mammalian species (Burrows, 1996; Parker, 2003; Poulet and Hedwig, 2006; Roberts et al., 2010).

Mapping the synaptic properties and monosynaptic connectivity rates in neocortex has been a central aim of in vitro slice studies, with visually guided multiple whole-cell patch clamp V_m_ recordings being the method of choice (Debanne et al., 2008; Deuchars and Thomson, 1995; Edwards et al., 1989; Feldmeyer and Radnikow, 2016; Feldmeyer et al., 2006; Geiger et al., 1997; Lalanne et al., 2016; Lefort et al., 2009; Markram et al., 1997b; Mason et al., 1991; Wang et al., 2015; Wozny and Williams, 2011; Yassin et al., 2010). The whole-cell recording technique has been adapted for use in vivo (Lee and Brecht, 2018; Margrie et al., 2002; Petersen, 2017) with more recent studies using dual, blind, whole-cell recordings to assess correlations of sub-and supra-threshold V_m_ activity between pairs of cortical neurons in awake mice (Arroyo et al., 2018; Gentet et al., 2010; Poulet and Petersen, 2008; Zhao et al., 2016). Dual V_m_ recordings provide a technical basis for testing for monosynaptic connectivity, but the likelihood of two cortical neurons being connected is low, dependent on cell type and negatively correlated with inter-somatic distance (Holmgren et al., 2003; Perin et al., 2011). Therefore, to identify connected pairs of neurons in vivo, it would help to be able to record from nearby, genetically labelled neurons using visual control.

Here we describe in detail an approach using in vivo two-photon microscopy to target whole-cell recordings to neighboring, fluorescently labelled layer 2/3 cortical neurons. We show that this technique can be used to evoke action potentials and isolate unitary excitatory and inhibitory postsynaptic potentials in postsynaptic neurons (Ferrarese et al., 2018; Jouhanneau et al., 2015; 2018). A troubleshooting table is provided (**Table 1**) and we go on to discuss potential improvements and future applications of this technique in assessing the link between monosynaptic transmission and cortical function.

**Table 1:** Troubleshooting during multiple whole cell patching

| Step | Problem | Possible reason | Solution |
| --- | --- | --- | --- |
| Surgery | Edema | Brain surface damaged | Take care when removing the dura as damaging the pia will result in tissue swelling. |
| | | Craniotomy too large | Keep size $\sim 700\mu\text{m}^2$ |
|  |  | Anesthesia | Isoflurane increases plasma volume which can induce swelling (Hamada et al., 1993). Drill a smaller craniotomy and use 1.2% agarose in Ringer's solution on the craniotomy to damp the swelling. Urethane does not increase plasma volume as much. |
| 2 | Pipettes unable to enter the brain | Dura intact | Re-attempt to remove the dura. |
|  |  | Blood vessel in the way | Make sure your brain entry point is clear. The use of green light will help to increase the contrast between blood vessels and brain tissue. |
| 3 | Dye not flowing out the pipette | Pipette clogged | Make sure the positive pressure is on before entering the ringer solution this will help maintaining a clean pipette tip. Use a fresh pipette. |
|  |  | Debris accumulating outside the pipette tip | Precipitate accumulating outside of the pipette could be a sign of grounding issue. Make sure the Ringer's solution in the recording chamber is not touching the head post. |
| | | Pipette clogged, visible debris inside the pipette | Debris inside the pipette can come from the intracellular solution itself. Use a fresh $0.45\ \mu\text{m}$ syringe filter (Minisart SRP4, Sartorius Stedium) for each experiment.<br>Debris can also come from the silver chloride electrode. Make sure the end of the pipettes are flame polished to avoid removing pieces of silver chloride coating.<br>Try clearing the tip of the pipette by increasing briefly the pressure (+ 50 mbar). If unsuccessful use new pipettes.<br>Do not use clogged pipettes even if the tip resistance is acceptable as it will most likely impair sealing. |
|  |  | Faulty pressure system | Check air pressure system can maintain a stable pressure. |
|  | No image | Laser off | Turn laser on. |
|  |  | Shutter closed | Open shutter. |
|  |  | PMT overload | Check external lights are switched off while the PMTs are on. Reset the PMT. |
|  | Poor imaging quality | High background fluorescence | Decrease the internal pipette pressure to reduce efflux of intracellular solution. |
|  |  | Leak of Ringer's solution out of the recording chamber | Check the contact between Ringer's solution and the objective. Try fixing the leak with Vaseline. |
|  |  | Laser power is to high | Small spherical dark spots appearing in the image is a sign of tissue damage caused by high laser power. The quality of the preparation is compromised and experiment should be terminated. |
| 4 | Unable to seal | Pipette resistance not optimal | Although lower resistance pipette ($< 5\ \text{M}\Omega$) will give you a better access to the cell it can also decrease sealing success. Aim for pipette resistance of $5 - 8\ \text{M}\Omega$. |
|  |  | Dirt on pipette tip | Although the resistance of the dirty pipette tip might be in the expected range, visible dirt dramatically reduce chances to seal successfully on neurons. Use a fresh pipette. |
| | | High pressure in Steps3/4 | Decrease the pressure to $< 30\ \text{mBar}$ while approaching the cell. Higher pressure will tend to "push" away the targeted neuron. In addition, in some cases, a slight negative pressure while sealing on the cell might be beneficial. |
|  |  | Intracellular solution | Check the osmolarity of your internal solution which usually need to be lower than the one of the Ringer's solution. |
| | | Holding potential not set to $-70\ \text{mV}$ | Make sure the holding potential is set up to $-70\ \text{mV}$ while sealing on the cell. In some cases, it will help to bring gradually the cell to $-100\ \text{mV}$ during the sealing procedure and then back to $-70\ \text{mV}$ before breaking in. |
| 5 | Unable to break in | Pipette resistance is too high | Pipette with a resistance higher than 8MΩ will tend to be more difficult to break in. The optimum pipette tip resistance is between 5 to 8 MΩ. |
|  |  | Patched a blood vessel | Blood vessels can look like cell bodies but a fast vertical scan will usually help identify cells from capillaries. Use a fresh pipette. |
|  |  | Patched on bundle of fibres | In some cases, the pipette might catch on fibres on the way to the cells of interest and even though a giga-seal will be made breaking in will fail. Use a fresh pipette. |
|  |  | Faulty pressure system | Make sure the pressure system is reactive to your suction. Suction must be brief. If something is damping the change of pressure breaking in will be impaired. |
| Recording | Short duration recordings (<5 mins) | Brain movement | Breathing of the mouse might create movement. Check the position of the mouse head relative to the body. If movement persists use 1.2% agarose ring solution to stabilize the brain movements, or stop the experiment.<br>Multiple pipettes entering the brain can cause a pressure build up in the surrounding tissue and the tissue will eventually relax to its original position. This may create tension on the seal and sometimes cause the pipette to push through or away from the cell. Visual checking during the recording using two-photon scanning and small adjustments of the pipette position can help stabilize recordings.<br>Isoflurane induces stronger pulsations of the brain than urethane. |
|  |  | Craniotomy is too large | New experiment with smaller craniotomy. If attempting awake recordings reduce craniotomy size even further. |
|  |  | Unstable head implant | The head implant may have become loosened due to tissue regrowth or poor gluing. Attempt adding extra glue or new experiment required. |
|  |  | Location of the pipette relative to the cell of interest | Aim for the most dorsal third of the targeted neuron soma to increase success rate and stability. |
| | Unable to trigger action potentials | Patched on glial cell | Check firing pattern, glial cells do not spike and typically have a very hyperpolarized $V_m$ with little or no spontaneous input. Change pipette and start over. |
|  |  | Access resistance is too high | Transiently applying negative pressure to the pipette tip. Slightly increase positive pressure during the final targeting approach. Improve brain stabilization procedure to reduce movement which can increase the access resistance during the recording. Retract and use a lower resistance pipette. |
| | $V_m$ depolarized | Recording solution | Use fresh Ringer's and intracellular solutions and check osmolarity. |
| | $V_m$ drift | Reference electrodes | Change or re-chloridize the recording and reference electrodes. |
|  |  | Metal head implant touching Ringer's solution | Isolate head implant from Ringer's solution in recording chamber. |
|  | No spontaneous activity | Anaesthesia level too high | Reduce isoflurane levels. |
|  |  | Body temperature is too low | Adjust the temperature controller. |

## MATERIALS AND METHODS

The aim of this article is to provide a description of multiple, 2-photon targeted whole-cell patch-clamp recordings to monitor monosynaptic connectivity in vivo. The procedure is described for an acute one-day experiment in anaesthetized mice. All experiments were performed according to protocols approved by the Berlin Animal Ethics committee (Landesamt für Gesundheit und Sociales, LAGeSo) and comply with the European animal welfare law.

### Two-photon microscope

In vivo two-photon microscopy with galvanometric scanning (Femto2D, Femtonics) is used to visualize neurons and the whole-cell recording pipettes (**Figure 1**). The microscope is fixed to an air damped table (Tuned damping table RS 2000, Newport). While our microscope can only move in the vertical Z axis, the experimental equipment, including pipette manipulators and headstages and mouse holder, are mounted on a shifting table (V380FM-L, Luigs and Neumann) allowing horizontal movements in X and Y. Two, photomultiplier tubes (PMTs) (GasAsP detectors, Hamamatsu) are used to detect light, one fitted with a 498 – 570 nm band pass filter and the second with a 598 – 700 nm band pass filter to enable detection of green and red fluorophores respectively. A CCD camera is coupled to the microscope and used at the start of the experiment to place the electrodes over the region of interest using a 4X objective (UPLFLN 4X, NA 0.13, W.D 17 mm, Olympus). Subsequently a 40X water immersion objective with a long working distance (LUMPLFLN 40XW, NA 0.8, W.D 3.3 mm, Olympus) is used to target cell soma of interest in a field of view of 200 x 200 µm (0.84 µm per pixel). The tunable (680 – 1080 nm), mode-locked Ti:Sapphire laser (Chameleon Ultra II, Coherent) is used to excite a wide range of fluorophores (e.g. GFP, Alexa 488, Alexa 594, tdTomato). The Pockel cell (E.O. Modulator, Conoptics) enables a fine control of the laser beam intensity. To avoid tissue damage during cellular two-photon imaging, we kept the laser power < 20 mW under the objective as we observed that damage can occur while targeting neurons with a laser power > 20 mW. The microscope is controlled by a Matlab (mathworks) based imaging data acquisition software (MES v4.0 software, Femtonics). Different combinations of pipette and cellular fluorophores can be used. For example, we used the red fluorophore Alexa 594 in the intracellular solution when using mice lines expressing GFP in neurons. Note that because of their different excitation spectra it is possible to use the red fluorophore Alexa 594 in the intracellular solution visible at 820 nm to target neurons expressing td-Tomato which are visible at 950 nm (Jouhanneau et al., 2018). For deeper recordings soma-restricted expression of fluorescent proteins may help improve depth resolution by reducing neuropil fluorescence (Baker et al., 2016).

**Figure 1.**
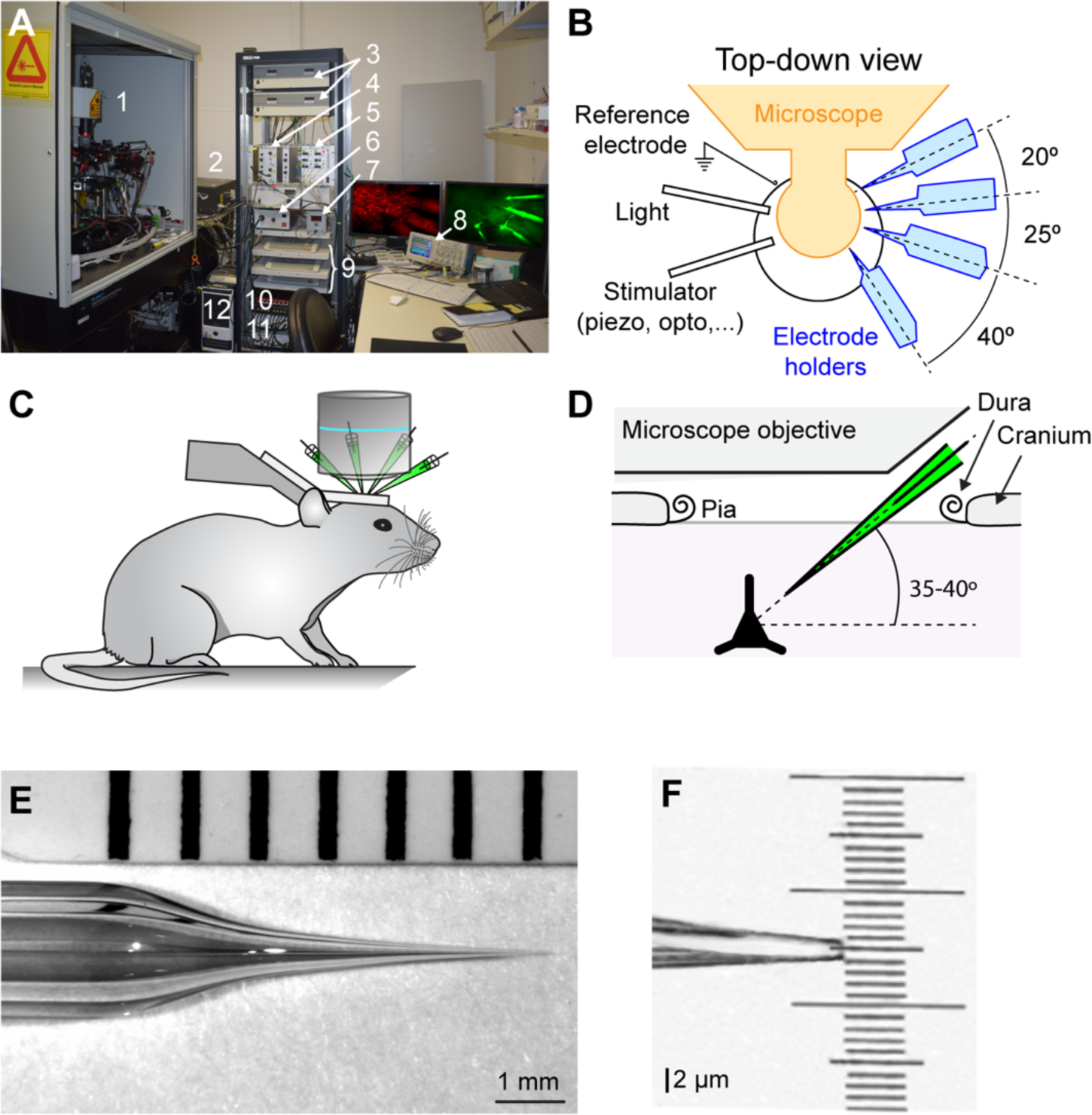
In vivo two-photon targeted multiple whole-cell patch-clamp setup. **(A)** Photograph of the setup showing (1) Two-photon microscope, micromanipulators and pre-amplifier fixed on an air-damped table; (2) Chameleon Ultra II laser; (3) Luigs and Neumann micromanipulator control units; (4): Sigmann Electronik air pressure controller; (5) Sigmann Electronik dual piezo amplifier (6) Light source to illuminate the preparation, (7) FHC temperature controller for anesthetized experiments, (8) Tektronix Oscilloscope, (9) Luigs and Neumman micromanipulator and shifting table control pads, (10) ITC-18 Heka analogue to digital converter board; (11) Multiclamp 700B amplifier; (12) Data acquisition computer. **(B)** Schematic top-down view of recording area showing arrangement of electrode holders, light, reference electrode and somatosensory/optogenetic stimulator. Note that all the recording electrodes are on the same side for easy of targeting and to allow space on the contralateral side for stimulation devices. **(C)** Cartoon showing mouse position and head support. **(D)** Schematic showing the angle of pipettes defined by the X-axis used to allow access under the objective. **(E)** Photograph of the glass recording pipette showing optimal taper for in vivo two-photon targeted patch-clamp recording. **(F)** Photograph showing a zoom of the 2 µm pipette tip from (E).

### Mice

The technique works with both wild type mice and strains expressing fluorescent proteins in subsets of neurons. In this article, we used mice aged between P18 – P30 from C57bl6J, NEX-cre (Goebbels et al., 2006) x Ai9 (Madisen et al., 2010), fosGFP (Barth et al., 2004), GAD67-GFP (Tamamaki et al., 2003), PV-cre (Hippenmeyer et al., 2005) x Ai9, or SST-cre (Taniguchi et al., 2011) x Ai9 background. Mice were maintained on a 12h light – dark cycle and had food and water *ad libitum*.

### Surgery

To expose the brain for recordings, mice are first anesthetized with 1.5% isoflurane and show an absence of tail pinch reflex and whisker movements. Eye ointment (Bei trockenen Auge, Visidic) is used to protect the eyes and body temperature is maintained using a closed loop system with rectal probe and heating pad (DC Temperature controller, FHC). All tools are cleaned and dry heat sterilized using hot glass beads sterilizer (Steri 250, Keller, Fine Science Tools) prior to surgery. The head is shaved and skull cleaned, if necessary, intrinsic optical imaging through the skull can be performed at this stage if functional localization of recording site is required. The connective tissue is carefully removed using forceps and micro-scissors (Fine science Tools) and the skull is cleaned using a microcurette (Fine science Tools) to remove any remaining tissue on the surface of the bone. A solution of 3% hydrogen peroxide (H_2_O_2_) can be applied for 30 seconds at this stage to help clean the exposed bone surface, however the bone can also be cleaned by gently scraping the bone with a razor blade. Next the skull is washed thoroughly with Ringer’s solution (135 NaCl, 5 KCl, 5 HEPES, 1.8 CaCl2, and 1 MgCl2135 NaCl, 5 KCl, 5 HEPES, 1.8 CaCl2, and 1 MgCl2) and thoroughly dried. Avoiding the recording site, the exposed skull is then lightly scratched with a 25G syringe needle to create grooves in the skull. It is important to remove any remaining hairs or dirt at this stage to avoid possible infection. Next, superglue (Loctite 401) is applied first at the edges of the exposed skull, to glue the skin to the bone, and then to the entire exposed skull surface avoiding the recording site. A lightweight metal head implant is then placed on the hemisphere contralateral to the recording site and covered with glue. To secure the head implant, dental cement is applied on top of the entire layer of glue. As the dental cement viscosity increases, a recording chamber with access for the recording pipettes can be modelled around the area of interest using a spatula. It is important to completely cover the head implant with glue and dental cement to avoid any possible contact between the Ringer’s solution and the metal of the head implant during the recording which can lead to electrical noise and voltage offsets.

Once hardened, the recording chamber is filled with Ringer’s solution. After a few minutes, the skull will become translucent and the blood vessels visible. Then, the Ringer’s solution should be removed to let the bone dry and a 500 µm diameter dental drill head (Komet, Brassler) operated by a dental drill (Success 40, Osada) is used to thin the skull over the recording site. The ideal craniotomy size is ∼ 700 µm^2^ for anesthetized mice; note that a craniotomy exceeding 1 mm in diameter will impair recording stability (see **Table 1**). Drilling is stopped as soon as blood vessels become clearly visible through the bone. This corresponds to a bone thickness of ∼ 100 µm (Papadopoulos et al., 2017). Bone dust is removed with wet tissue paper and the chamber is refilled with Ringer’s solution. A 30G syringe needle is used to pick away the final layer of bone with great care. Next, a durectomy is made using a smaller diameter needle (e.g. 29G), with a handmade small hook at the tip of the needle. Adjusting the angle of illumination of the craniotomy is key to visualizing the dura (∼ 30°). The handmade hooked-tip of the needle is used to gently lift the dura away from the future spot of pipette insertion.

### Whole-cell pipettes and electrophysiological equipment

We use a four-step pulling custom program on a Sutter puller (Model P-1000, Sutter instrument) with 2 mm diameter borosilicate capillaries (Hilgenberg) to pull 5 – 8 MΩ pipettes. The first two steps of the pulling program are identical and used to create a taper of ∼ 6 mm, the third step is short and design to decrease the diameter of the capillary, and finally the fourth step is used to create a tip of ∼ 2µm (**Figure 1E, F**). The taper is slightly longer than that typically used in vitro to avoid causing excess pressure on surrounding tissue and possible damage. Three to four pipettes are filled with intracellular solution containing, in mM: 135 potassium gluconate, 4 KCl, 10 HEPES, 10 phosphocreatine, 4 MgATP, 0.3 Na3GTP (adjusted to pH 7.3 with KOH), 25 μM Alexa-594 (Invitrogen) and 2 mg/ml biocytin. Pipettes are next fixed to a pipette holder (Molecular Devices) mounted on a LN Junior 3-axis (X, Y, and Z) micromanipulator with low drift and a long traverse path (up to 22 mm on the X axis) where the X axis is angled at 35 – 40° (**Figure 1D**) (Luigs and Neumann). An Ag/AgCl ground electrode is next placed into the recording chamber filled with Ringer’s solution and electrophysiological signals are amplified using Axon Instruments amplifiers Multiclamp 700B (Molecular Devices). The analog signals recorded are filtered at 10 kHz and digitized at 20 kHz using the analog/digital converter ITC-18 board (Heka) and IgorPro (Wavemetrics) running on a Windows PC. For online visualization of the electrophysiological signal we use an oscilloscope (Tektronix TDS2024C). To allow easier and faster access to the exposed brain and space for stimulators on the contralateral side, all pipettes are positioned on one side of the preparation (**Figure 1B**).

### Multiple two-photon targeted whole-cell patch clamp recordings

As soon as the pipettes are inserted into the pipette holders, a 180 – 200 mbar positive pressure is applied via a syringe. A manual-seal-sucker (Sigmann Elektronik GmbH) manometer is used to monitor the pressure applied to all channels independently. All electrodes are moved into the Ringer’s solution in the recording chamber in voltage-clamp seal-test mode to monitor the pipette tip resistance on the oscilloscope.

#### Step 1: Positioning above brain (Figure 2A)

Using the low magnification 4X objective with green light illumination and the CCD camera, the pipettes are placed under positive pressure (∼ 200 mbar) into the Ringer’s solution and then the tips are moved to within ∼ 20 – 30 µm apart from each other and ∼300 µm above the craniotomy. At this time, the pipette resistance is checked (5 – 8 MΩ; see Table 1). Then, by switching to the higher magnification 40X objective the Ringer’s solution comes in contact with the objective and the pipettes tips are moved to the same focal plan. The coordinates of the pipette micromanipulator control pads are reset to zero. Next, we use the experimental stage micromanipulators (X and Y axis) and the objective focus (Z axis) to inspect the brain surface and find an entry point clear from large blood vessels, dirt or irregular surfaces. Clean entrance of the pipettes into the brain is critical for successful patching. The coordinates of the manipulator units controlling the stage are noted at the selected insertion point as a reference to help guide the movement of the pipette tips onto the brain surface. Next, the focus is moved back up to the pipette tips which are then carefully aligned. The focal plan is moved to the brain surface and the pipettes are lowered vertically one by one using a medium control sensitivity of the micromanipulator control pads (28 µm per handwheel rotation). As the pipettes are lowered, slight lateral movements are performed to help visualize the shadow of the tips. Because of the positive pressure applied to the pipette, as soon as the pipette gets into contact with the brain a clear depression can be seen on its surface which will coincide with a sudden increase in resistance of about ∼ 20% of the peak value (as observed by a decrease in the current step amplitude on the oscilloscope). At this point, the pipette micromanipulator control pad values are reset to zero.

**Figure 2.**
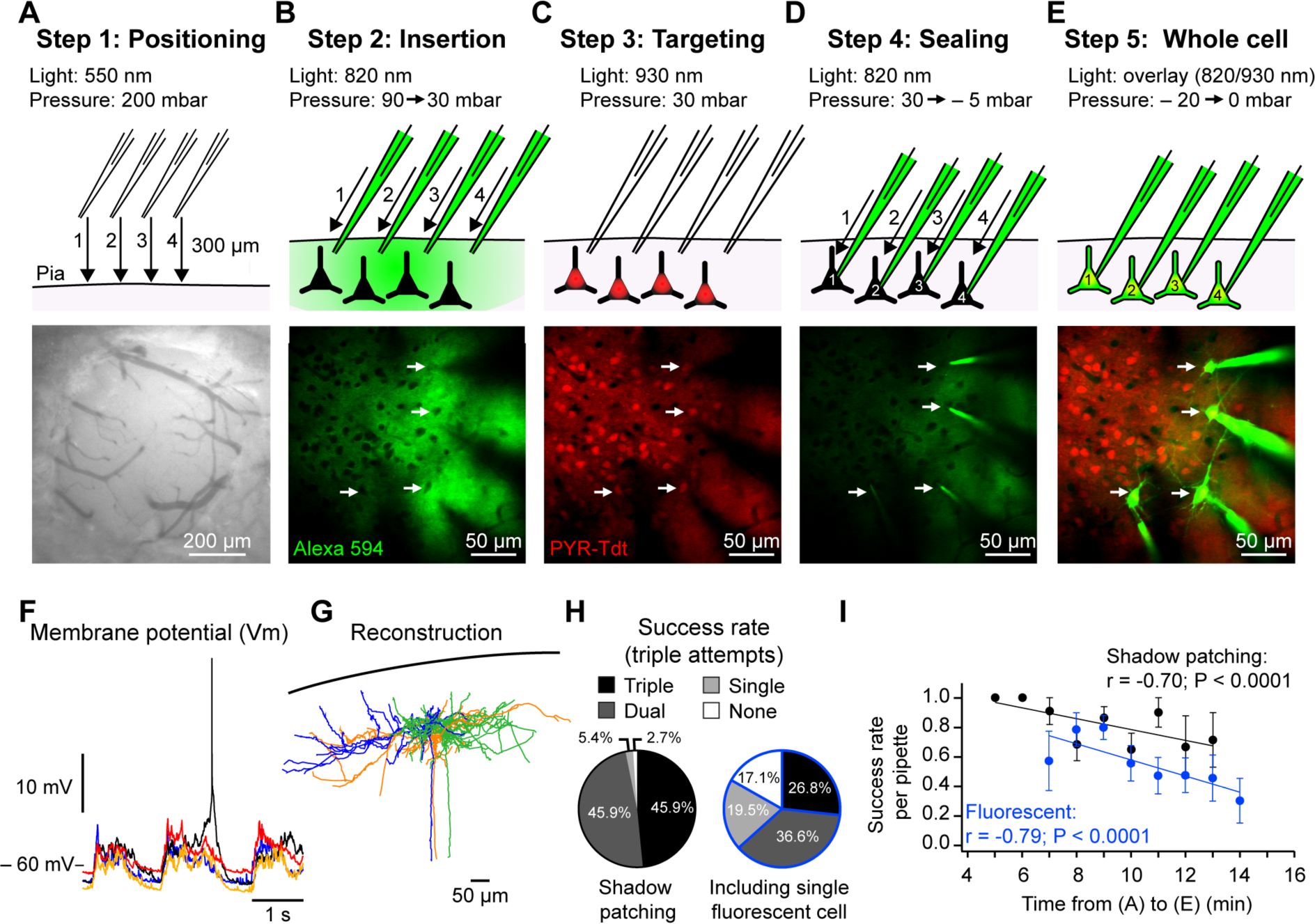
Two-photon targeted whole-cell patch-clamp procedure. **(A)** Step 1. Top: Schematic showing position, movement direction (arrow) and movement order (1 – 4) of dye-filled (Alexa 594) recording pipettes from ∼ 300 µm above the craniotomy to the brain surface, high positive pressure is maintained to avoid pipette tip clogging. Bottom: photograph taken using a CCD camera illuminated with green light showing example craniotomy used for anesthetized patching (∼ 500 µm diameter) in which the dura has been removed. **(B)** Step 2. As in (A), but showing, top: the insertion of the pipettes through the pia under visual control (820 nm) using high positive pressure to ∼ – 100 µm from pial surface. As soon as the pipettes enter the brain the pressure is reduced to 100 mbar and then 30 mbar when closer to the cell body positions. Bottom: In vivo two-photon image showing the position of 4 pipettes in same focal plan near the cell bodies of interest. Positioning is performed sequentially, one pipette at a time. Cell bodies are revealed as dark shadows contrasting with the fluorescent signal of the extracellular dye. **(C)** Step 3. Top: the targeting phase where the excitation light wavelength is altered to visualize the cells of interest; in this case, excitatory glutamatergic neurons expressing the red fluorophore tdT (white arrows). Bottom: in vivo fluorescent image of pyramidal neurons expressing tdT corresponding to the photo in (B). **(D)** Step 4. Top: the final sealing phase of the procedure. A recording pipette is pushed into the cell soma membrane and then, upon strong contact, the pressure is released to achieve a giga-seal. The cells are sealed sequentially under visual control. Bottom: in vivo image following sealing of all 4 pipettes. Note the reduction in background fluorescence during sealing because of the reduction in extracellular dye. **(E)** Step 5. Top: entering whole-cell configuration following seal-breaking by applying a transient negative pressure. As soon as the membrane patch is ruptured the dye within the pipettes will fill the neurons revealing anatomy. Bottom: in vivo image of neurons filled with Alexa 594 after the recording experiments. **(F)** Simultaneous example in vivo whole-cell V_m_ recordings of the four excitatory pyramidal neurons shown in (E) showing spontaneous activity with Up-and Down-states under urethane anesthesia. **(G)** Post-hoc reconstruction of three biocytin-filled excitatory pyramidal neurons from a multiple whole-cell recording. **(H)** Proportion of triple, double, single and no recordings from trials using three pipettes in (left) wild type mice using the shadow patching method and (right) in mice expressing a cell type specific fluorophore (PV-cre x Ai9 and SST-cre x Ai9). Data from fluorescent mice included at least 1 fluorescent neuron in the single/double/triple recordings. **(I)** Plots showing the negative correlation between the success rate of achieving a whole-cell patch clamp recording in wild type and fluorescent mice (same data as in (H)) and the time taken from phases (A) to (E) described above. Each dot represents the mean success rate for a 1 min time bin from 15 wild type mice (37 trials) and 18 mice expressing fluorescent proteins (SST-cre x Ai9 and PV-crex Ai9; 42 trials).

#### Step 2: Entering the brain (Figure 2B)

Using the highest sensitivity speed on the micromanipulator (3 µm per handwheel rotation), the pipettes are slowly moved through the pia one-by-one. During insertion into the brain, the pipette resistance will gradually increase and then suddenly return to their initial value as they break throughout the pial surface. Then the pressure is reduced to 70 – 90 mbar. Next, two-photon imaging is used to move the pipette tips to – 50 µm depth using the X-axis focus. Because of the positive pressure, the dye (Alexa-594) contained in the intracellular solution will diffuse into the neuropil and highlight blood vessels, cell soma and dendrites as dark “shadows” allowing targeting cells of interest even in wild type mice (Kitamura et al., 2008). Care should be taken during this step not to use too high laser power as it may cause tissue damage (see **Table 1**).

#### Step 3: Targeting cells of interest (Figure 2C)

Having lowered the pressure to 70 – 90 mbar, the pipettes are then moved one-by-one to a depth of – 150 to – 200 µm (border of cortical layers 1 and 2) using the highest sensitivity movement setting. During pipette travel through the brain, great care is taken to avoid cells bodies and capillaries using both visual control from the two-photon imaging and the seal test pulse on the oscilloscope. In wild-type mice, without expression of fluorescent proteins, the contrast between the dye in the neuropil and the dark unlabeled cells, the shadow patching technique (Kitamura et al., 2008), can be used to target neurons of interest. With experience, the dendritic shape of the cell can help identify excitatory from inhibitory neurons. Lateral movement should be kept to a minimum with a maximum of 100 µm per pipette. Then the pressure is decreased to 30 mbar and the pipette micromanipulator controls are switched to a stepping mode (2 µm per step).

#### Step 4: Sealing (Figure 2D)

Next, the final approach and seal is performed sequentially, one pipette at a time. The pipette voltage offset is set to 0 mV and the first pipette tip is lowered using 2 µm steps onto the cell membrane using the X-axis while watching the oscilloscope closely to observe changes in tip resistance. The sign of a good contact between the pipette tip and the neuron membrane is when the seal test pulse reduces in amplitude and fluctuates with a wave like pattern. In contrast, an abrupt and sustained reduction in pulse amplitude (i.e. resistance increase) without fluctuations is typical of a contact with a capillary. Good contact can sometimes be visualized during two-photon scanning as a small, expanding, fluorescent dimple in the cell membrane. As soon as a good first contact has been observed, one or two further steps are made and the positive pressure immediately released followed by a transient negative pressure to optimize the seal. V_m_ holding voltage is immediately placed at – 70 mV to help improve seal. Typically, this leads to a large reduction in the amplitude of the test pulse and the formation of a giga-seal, however light manual suction is sometimes required to improve the quality of the seal and/or transiently hyperpolarizing the cell to – 100 mV. This procedure is then repeated with the other pipettes one after the other.

#### Step 5: Whole-cell configuration (Figure 2E)

When all the pipettes are sealed onto the targeted neurons, a brief and gentle suction is used to break the membrane and enter whole-cell configuration on each of the neurons with a giga-seal. With the whole-cell configuration established, we next slowly retract each pipette away from the cell body ∼ 5 µm using the X axis used for the final approach to the cell. All recordings are then switched to current clamp mode for V_m_ recordings.

### Intracellular current injection

After allowing the cells to recover (∼ 2 – 3 min), we next use intracellular current stimulation protocol to characterize their intrinsic properties. In our experiments, each neuron receives 500 ms square current injections ranging in amplitude of – 200, – 100 pA, and then 50, 100, 150, 200 pA. This helps define rheobase, intrinsic excitability, and firing pattern of the recorded neurons. Next, hyperpolarizing current pulses of – 100 pA, 200 ms duration, 200 ms interval, are applied for 30 s to determine the input resistance followed by 30 s without stimulation to record spontaneous sub-and supra-threshold activity. The access resistance should be < 50 MΩ, high access resistance can make it difficult to inject sufficient current to evoke single spikes, filters action potential recordings and makes estimation of the V_m_ during current injection problematic. Next, we define the square current pulse amplitude necessary to drive the recorded neuron to fire a single action potential. We aim to use the smallest duration possible, usually between 10 to 15 ms of 100 to 400 pA amplitude, however higher amplitude and shorter duration pulses could be attempted. After establishing these parameters, we stimulated at 0.5 or 1 Hz (**Figure 3C**).

**Figure 3:**
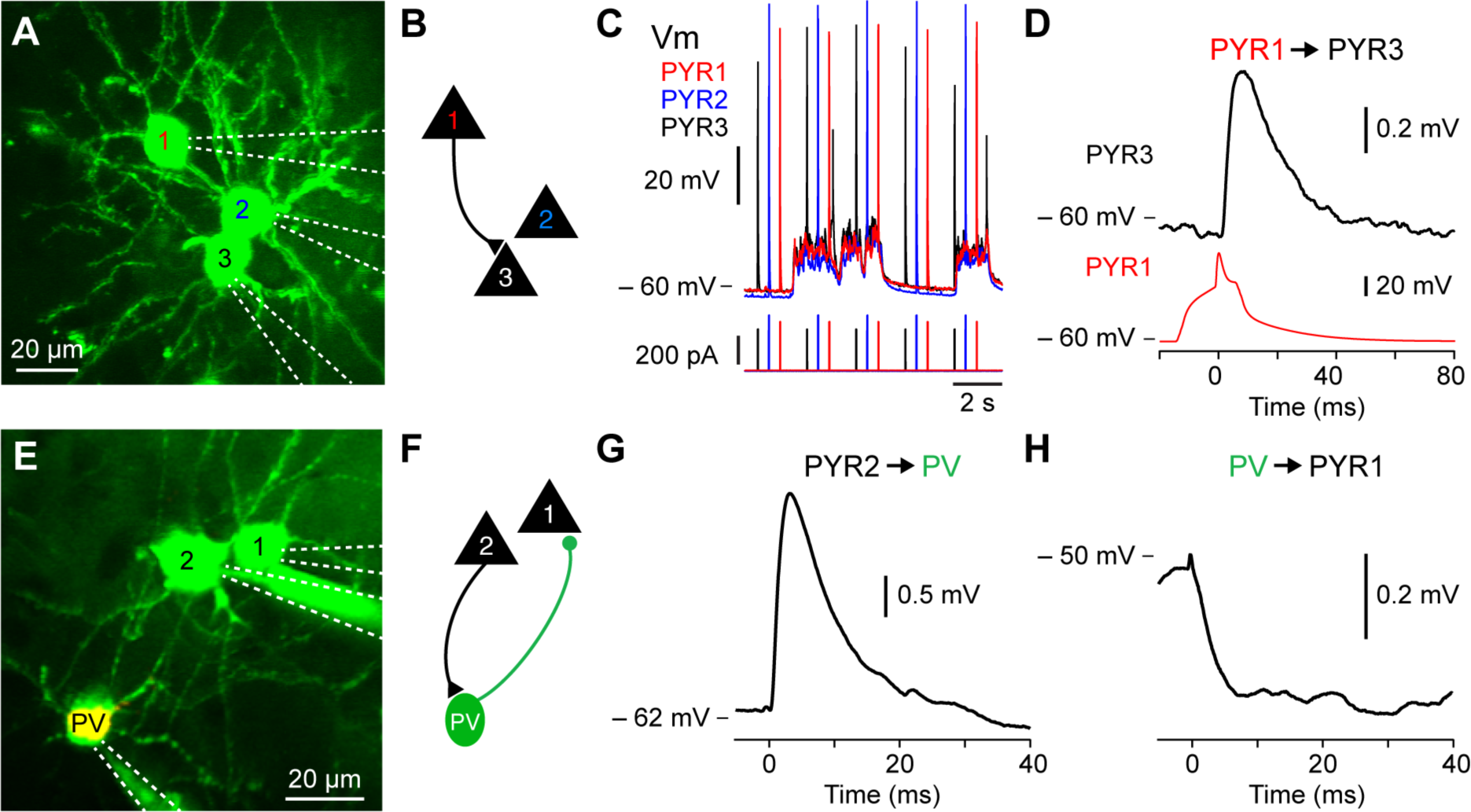
Excitatory and inhibitory monosynaptic connections in vivo. **(A)** In vivo two-photon fluorescent imaging of 3 excitatory pyramidal neurons filled with Alexa 594. **(B)** Connectivity diagram of experiment shown in (A) where PYR1 is connected to PYR3. **(C)** V_m_ fluctuation of the pyramidal neurons recoded in (A) showing the stimulation protocol consisting of brief current injection to evoke single action potentials in each neuron every 2 seconds. **(D)** Example monosynaptic excitatory connection from PYR1 to PYR3 shown in (A-C). **(E)** In vivo image of a triple recording including a PV-tdT expressing GABA-ergic inhibitory interneuron. **(F)** Connectivity diagram of experiment from the triple recording shown in (E) where an excitatory pyramidal neuron PYR2 is connected to PV-tdT expressing GABA-ergic inhibitory PV, while PV sends an inhibitory monosynaptic connection to the excitatory pyramidal neuron PYR1. **(G)** Excitatory monosynaptic connection from PYR2 to PV shown in (E,F). **(H)** Inhibitory monosynaptic connection from PV to PYR1 shown in (D).

There was no tonic current injection applied during the recording to avoid misestimation of the V_m_ due to possible changes in the access resistance during the recording. Recordings are terminated when the most hyperpolarized sections of the Downstate V_m_ are more depolarized than – 50 mV. Due to differences in ionic concentration, valency and mobility between the intracellular and extracellular solution, a Liquid Junction Potential (LJP) will be established when the pipette enters the recording chamber (Barry and Lynch, 1991). The LJP can be ∼ 10 mV and is complex to calculate accurately in vivo, therefore, to avoid miscalculation, we do not subtract the LJP from the recorded values.

### Identifying a connection

In anaesthetized Downstates or during hyperpolarized phases of network activity in awake animals, even small amplitude (0.1 – 0.5 mV) monosynaptic connections can typically be observed by eye in single trials. However, online, running averages of the postsynaptic response to an evoked spike helps monitor the presence of a connection as well as the quality of the recording. To confirm the presence of a connection post hoc, we used a non-parametric two-tailed Wilcoxon signed rank test comparing trial-by-trial amplitude measurements of the connection with shuffled measurements. We also used a bootstrapping method in which we compared a randomly selected, with replacement, amplitude measurements from the individual trial responses with those from shuffled measurements of amplitude. Next, we calculated the mean response amplitude and the mean shuffled, noise amplitude from the bootstrapped distributions. To obtain the 95% confidence intervals we then repeated this process 10,000 times (see Jouhanneau et al., 2015).

### Anatomy: live fluorescent two-photon imaging and post hoc biocytin staining

During a successful recording, the fluorophore Alexa 594 diffuses into the neurons and allows live visualization of the cell’s anatomy (**Figures 2E, 3A and 3E**). Stacks of scans at 820 nm wavelength separated by 2 µm can help identify the cell type using the somatic and dendritic anatomy as well as the presence of dendritic spines. However, for higher resolution anatomical reconstruction mice are deeply anaesthetized with an i.p. injection of urethane (2.5 g/kg, Sigma-Aldrich) before being transcardially perfused with cold Ringer’s solution and then by 4% paraformaldehyde solution (PFA, Roti-Histofix 4%, Roth). After perfusion, the brain is removed and placed in 4% PFA overnight at 4°C and then in phosphate buffer (Roti-CELL 10×PBS, Roth) and stored at 4°C until further processing.

Tangential slices of 100 µm are cut using a Leica VT1000 S vibratome and stored in phosphate buffer. Intracellular staining with biocytin is then revealed using a standard ABC kit (Vectastain Elite ABC-Peroxidase kit, Biozol), with diaminobenzidine (DAB, Vector lab) enhancement. Treated slices are mounted on glass slides using a gel mounting agent (Moviol, Sigma-Aldrich), sealed with nail polish and stored at 4°C. Reconstructions of the recorded neurons are performed using the software NeuroLucida (MicroBrightField) **(Figure 2G)**.

### Success rates

We next calculated the success rates of our approach during patching of layer 2 neurons (depth: −182.0 +/− 2.5 µm; distance: 39.0 +/− 1.8 µm) in 37 trials, each trial corresponding to one insertion of 3 pipettes into the brain, in 15 anesthetized wild-type mice (males, 22.0 +/− 0.3 days old) using the shadow patch method performed by a trained researcher. We calculated the number of times we were unsuccessful or obtained a single, dual or triple whole-cell recording. In 17/37 trials we obtained a triple recording, in 17/37 trials a dual, in 2/37 single and in 1/37 no recordings (**Figure 2H**). Thus in 92% of shadow patching trials using three pipettes we obtained at least a dual recoding that would allow a connectivity test. We then repeated this analysis for attempted triple recordings (3 pipettes) in mice expressing a fluorescently labelled indicator in a subset of GABA-ergic interneurons (PV-tdT or SST-tdT), where a successful recording trial had to include a least one fluorescently labelled neuron. In 11/41 trials from 18 mice we obtained a triple recording, in 15/41 trials double, 8/41 single (ie one tdT labelled neuron recorded) and were unsuccessful in 7/41 attempts (**Figure 2H**). Thus, in 63 % of trials with fluorescently labelled mice we obtained at least a dual recording including 1 fluorescent GABA-ergic neuron to allow for a connectivity test.

During these experiments, we noticed that for trials that took longer it seemed harder to perform a successful whole-cell recording. From our triple recording dataset in Figure 2, we therefore systematically recorded the time to go from Step 1 (positioning above brain) to Step 5 (whole-cell recording). Plotting the time taken against the success rates of successfully patching one neuron showed a significant negative correlation and confirmed that reducing the time taken to patch improves success rates for patching (**Figure 2I**). In a different set of experiments where the recordings were not terminated prematurely and the Downstate V_m_ was < – 50 mV, we calculated a mean recording time of 15 +/- 6 minutes (n = 143 cells) with a minimum recording time of 5 minutes and a maximum of 32 minutes.

### In vivo glutamatergic excitatory monosynaptic inputs to excitatory pyramidal neurons and GABA-ergic inhibitory interneurons

During slow wave sleep and under anesthesia the V_m_ of cortical neurons fluctuates between hyperpolarized, synaptically quiescent, Downstates and depolarized, synaptically active, Upstates (Steriade et al., 1993). We first examined synaptic transmission between excitatory pyramidal neurons in Downstates. In wild type mice, pyramidal neurons were targeted using their pyramidal shaped soma and apical trunk visible as shadows against the fluorescent extracellular space and after each successful recording, confirmed using Z-stack images to visualize the somatic and dendritic morphology (e.g. spines). Moreover, we used transgenic mice expressing fluorophore in excitatory pyramidal neurons (PYR) using offspring of the NEX-cre line crossed with the Ai9 reporter mouse to study PYR to PYR monosynaptic connectivity. To trigger spikes and measure synaptic transmission we depolarized each neuron with injection of a short depolarizing current 100 – 400 pA of 20 – 50 ms duration at 0.5 or 1 Hz to evoked a single action potential (**Figure 3B**) and used spike triggered averages to look at the corresponding unitary excitatory postsynaptic potential (_u_EPSP). To study short term synaptic dynamics, multiple action potentials could be triggered by increasing the current duration number and time.

We went on to use the same approach to examine excitatory connections from PYRs to different subtypes of GABA-ergic neurons including parvalbumin (PV), somatostatin (SST) and vasoactive intestinal polypeptide (VIP) expressing neurons. The PV-cre, SST-cre and VIP-cre mice were crossed with the Ai9 reporter line to visualize the subpopulation of GABA-ergic neurons of interest. The approach to target recordings of interneurons is technically similar to that of excitatory neurons. However, in some cases, during the sealing step, the positive pressure was lower than usual (∼ 10 mbar) in order to target small diameter neurons like VIP interneurons. For further details on inputs to PV and SST interneurons in vivo see (Jouhanneau et al., 2018).

Using this approach, we found that barrel cortex layer 2 excitatory pyramidal neurons had a connectivity rate of 6.7% (Jouhanneau et al., 2015), while connections from excitatory pyramidal neuron to PV neurons was 44.4% and to SST neurons was 43.6% (Jouhanneau et al., 2018). For further details see (Jouhanneau et al., 2015) and (Jouhanneau et al., 2018).

### Inhibitory monosynaptic connections to PYRs and INTs in vivo

We next used multiple two-photon targeted patch-clamp recordings to examine inhibitory monosynaptic neurotransmission in vivo. Here we recorded inhibitory connections from GABA-ergic INTs to PYRs and also from INT to INT. The connectivity rate from layer 2 barrel PV neurons to excitatory neurons was 60.6% and from SST neurons to excitatory neurons 47.1% (see Jouhanneau et al., 2018). The recording procedure is similar to that described above, but because interneurons often have a higher input resistance, small amplitude and shorter duration current steps are used to trigger single action potentials. Moreover, because of their hyperpolarized reversal potential, _u_IPSPs are more visible at more depolarized postsynaptic V_m_ values. This was evident in our recordings, where the amplitude of _u_IPSPs was larger in Upstates compared to Downstates (see Jouhanneau et al., 2018).

## DISCUSSION

Understanding the link between monosynaptic connectivity and the functional properties of cortical neurons is a key goal of neuroscience. Here we have described an approach that allows V_m_ recordings of monosynaptically connected cortical neurons in vivo. The setup uses a standard in vivo two-photon microscope, whole-cell patch clamp amplifiers and motorized micromanipulators. With training, multiple whole-cell recordings of neurons in layer 2 can be performed with a success rate of forming a dual recording of ∼ 90% and recording duration (∼ 15 min) similar to single electrode, blind, in vivo patch clamp recordings. Perhaps the key indicator of patching success is an unhindered pipette entrance into the brain and rapid progress through the tissue (**Figure 2**). In **Table 1** we have outlined a list of common problems with targeted patch-clamp recordings and possible solutions. Here, we discuss the key features, limitations and future perspectives for multiple, targeted in vivo whole-cell recordings.

Increasing the number of pipettes per trial helps test more possible connections with 2 pipettes allow the testing of 2 possible connections, 3 allowing 6 tests and 4 12 tests. More pipettes provide an opportunity to not only to improve the changes of identifying a connection but also look at higher order connectivity motifs (Guzman et al., 2016; Peng et al., 2017). While we have successfully used 4 pipettes to obtain quadruple whole-cell recordings (**Figure 2A-E**), our data on success rates (**Figure 2H-I**) was taken from a series of experiments using 3 pipettes. In wildtype mice, during shadow patching, 4 pipettes or more could be a significant advantage to help increase the yield of recorded cells. However, for targeting fluorescently labelled subsets of neurons, the experimenter needs to weigh the advantage of using a 4^th^ pipette against the extra time taken to insert 4 pipettes into the brain and target the labelled neurons.

A limitation of our approach is the use of anesthesia during the recording session. While multiple whole-cell recordings have been performed in awake animals (Arroyo et al., 2018; Gentet et al., 2010; Poulet and Petersen, 2008; Zhao et al., 2016), little data exists on monosynaptic transmission in awake animals (Jouhanneau et al., 2018; Pala and Petersen, 2018). The increased movement of the brain in awake animals not only limits the chances of forming a seal between the pipette and the cell membrane but also reduces the recording duration preventing longer term plasticity protocols (e.g. spike timing dependent plasticity (Bell et al., 1997; Markram et al., 1997a). The use of agarose or glass cover slips on the brain surface has helped reduce movement during imaging experiments and can be used for targeted whole-cell recordings. Moreover, having the mouse standing on a trackball or platform with suspension can help reduce the pressure exerted on the head during leg movements. Together, these approaches may help stabilize the brain for longer duration recordings both in anesthetized and awake mice.

The approach presented above focusses on recordings from superficial layer cortical neurons. Moreover, as with the vast majority of cortical slice work, the neurons targeted were closely positioned (< 150 µm apart). It is important to examine synaptic transmission between deeper and more distant neurons, perhaps even in different cortical regions. Electrodes can easily be positioned to target different parts or depths of the brain, but both the depth and field of view are determined by the optical properties of the microscope. The combination of cell type specific mouse lines (Daigle et al., 2018; Gerfen et al., 2013; Harris et al., 2014) with improved depth resolution 2-photon microscopes (Papadopoulos et al., 2017) has provided optical access to granular and infragranular layers and may make multiple targeted recordings possible in deeper layers. Moreover, new two-photon microscope designs with larger fields of view could allow experimenters to examine neurons situated 1000s of micrometers apart (Sofroniew et al., 2016; Stirman et al., 2016). Even with improved microscopes, however, the scattered fluorescence from the extracellular dye puffed out during patching remains a problem for accurate visualization of the pipette tip and targeted recordings. One possibility may be to use a coating material on the pipette tip to limit light scatter and improve contrast of the tips (Andrasfalvy et al., 2014).

Cortical excitatory neurons are sparsely connected and therefore a key limitation to the throughput of any connectivity study is to find and record from connected pairs. Both in vitro and in vivo studies are normally made blind to connectivity which can lead to many frustrating recordings from unconnected neurons. One way to address this is to increase the numbers of recording pipettes to allow the testing of more connections per recording session. This has been successfully implemented in vitro (Peng et al., 2017; Perin et al., 2011), but will require more challenging surgery and manipulation of the pipettes in vivo. Another approach could be to use transsynaptic tracing to visualize connected pairs prior to recording (Wickersham et al., 2007). So far, however, single cell initiated transsynaptic tracing has been used with sequential rather than simultaneous anatomical tracing (Vélez-Fort et al., 2018), or calcium imaging (Wertz et al., 2015) of presynaptic neurons. With the development of less toxic rabies viruses variants (Ciabatti et al., 2017; Reardon et al., 2016), however, this approach could now be attempted with simultaneous recordings from pre-and post-synaptic neurons.

Our approach allows a limited number of cells (2 – 4) to be tested for putative connections, but cortical neurons integrate synaptic inputs from thousands of presynaptic neurons. To investigate synaptic integration further, it will be important to be able to activate unitary monosynaptic inputs from more than 1 neuron with high temporal precision. The recent development of in vivo single cell optogenetic stimulation (Packer et al., 2015; Rickgauer et al., 2014), has provided a way to activate multiple single neurons with high resolution spatial and temporal patterns. A combination of this technique with in vivo whole-cell recordings to monitor small amplitude subthreshold synaptic inputs, could provide an exciting method to investigate the integration of multiple unitary inputs in vivo.

An in vivo patch clamp recording session can be slow, especially when learning the technique or using multiple electrodes. In particular, the replacement of old pipettes with unused ones at each new recording attempt takes up valuable time. A recent study has circumvented this problem with the use of a commercially available detergent and rinsing procedure (Kolb et al., 2016). This allowed the reuse of the same pipettes with no degradation in signal fidelity both in vitro and in vivo. Robotic assistance to move the pipettes also may help speed up this process and has recently been implemented for the entire visualized patching process (Annecchino et al., 2017; Suk et al., 2017).

The whole-cell technique allows intracellular access to the recorded neurons and future work could make more detailed anatomical and genetic characterization of the recorded neurons. For example using single-cell RNA sequencing (Boldog et al., 2018; Cadwell et al., 2016; Fuzik et al., 2016; Jiang et al., 2015; Muñoz et al., 2017) or higher resolution bright field (Feldmeyer et al., 2006) or electron microscopic (Fernández et al., 1996) anatomical analysis of the recorded synaptic connections.

The craniotomy and glass recording pipette exposes the brain and requires the use of extracellular Ringer’s solution as well as an intracellular solution. These solutions are made in the lab and therefore provide an access point for the application of extra-and intra-cellular (Ferrarese et al., 2018; Palmer et al., 2014), pharmacological agents in vivo. For example, we recently applied intracellular blockers of different ion channels via the intracellular solution to investigate their impact on synaptic integration during network activity (Ferrarese et al., 2018), and extracellular antagonist to monitor the impact of acetylcholine on monosynaptic excitatory transmission between PYR neurons and neighboring SST GABA-ergic neurons (Urban-Ciecko et al., 2018). Future work could therefore use specific pharmacological agents to help understand the ionic mechanisms of synaptic transmission in active, intact networks.

## Conclusions

Two-photon targeted multiple-whole cell recordings provide a high resolution and cell-type specific way of identifying monosynaptically connected neurons *in vivo*. This approach will allow studies into the impact of network activity on synaptic transmission, the synaptic mechanisms underlying action potentials generation and link connectivity to functional responses at a millisecond time scale. Moreover, the possibility to record the V_m_ of both pre-and post-synaptic neurons provides a way to examine the synaptic basis of correlated spiking activity of cortical neurons.

## AUTHORS CONTRIBUTIONS

JSJ performed the experiments and analyzed the data. JSJ and JFAP designed the study and wrote the manuscript.

## ACKNOWLEDGMENTS

We would like to thank Janett König, Charlene Memler and Femtonics for technical assistance, Mario Carta, Sylvain Crochet, Alison Barth and Evgeny Bobrov for comments on earlier version of the manuscript. We thank Poulet lab members for help and constructive comments. This work was funded by the European Research Council (ERC-2015-CoG-682422, J.F.A.P.), the European Union (3×3Dimaging 323945), the Deutsche Forschungsgemeinschaft (DFG, FOR 1341, FOR 2143, J.F.A.P), the American National Institute of Health (NIH R01NS088958, J.F.A.P.), the Thyssen Foundation (J.F.A.P.), and the Helmholtz Society (J.F.A.P.).

## References

Andrasfalvy, B. K., Galiñanes, G. L., Huber, D., Barbic, M., Macklin, J. J., Susumu, K., et al. (2014). Quantum dot-based multiphoton fluorescent pipettes for targeted neuronal electrophysiology. Nat. Methods 11, 1237–1241.

Annecchino, L. A., Morris, A. R., Copeland, C. S., Agabi, O. E., Chadderton, P., and Schultz, S. R. (2017). Robotic Automation of In Vivo Two-Photon Targeted Whole-Cell Patch-Clamp Electrophysiology. Neuron 95, 1048–1055.e3.

Arroyo, S., Bennett, C., and Hestrin, S. (2018). Correlation of Synaptic Inputs in the Visual Cortex of Awake, Behaving Mice. Neuron 99, 1289–1301.e2.

Baker, C. A., Elyada, Y. M., Parra, A., and Bolton, M. M. L. (2016). Cellular resolution circuit mapping with temporal-focused excitation of soma-targeted channelrhodopsin. eLife 5, 11981.

Barry, P. H., and Lynch, J. W. (1991). Liquid junction potentials and small cell effects in patch-clamp analysis. J. Membr. Biol. 121, 101–117.

Barth, A. L., Gerkin, R. C., and Dean, K. L. (2004). Alteration of neuronal firing properties after in vivo experience in a FosGFP transgenic mouse. J. Neurosci. 24, 6466–6475.

Barthó, P., Hirase, H., Monconduit, L., Zugaro, M., Harris, K. D., and Buzsáki, G. (2004). Characterization of neocortical principal cells and interneurons by network interactions and extracellular features. J. Neurophysiol. 92, 600–608.

Bell, C. C., Han, V. Z., Sugawara, Y., and Grant, K. (1997). Synaptic plasticity in a cerebellum-like structure depends on temporal order. Nature. 387, 278–281.

Berry, M. S., and Pentreath, V. W. (1976). Criteria for distinguishing between monosynaptic and polysynaptic transmission. Brain Res. 105, 1–20.

Boldog, E., Bakken, T. E., Hodge, R. D., Novotny, M., Aevermann, B. D., Baka, J., et al. (2018). Transcriptomic and morphophysiological evidence for a specialized human cortical GABAergic cell type. Nat. Neurosci. 21, 1185–1195.

Bruno, R. M., and Sakmann, B. (2006). Cortex is driven by weak but synchronously active thalamocortical synapses. Science 312, 1622–1627.

Burrows, M. (1996). The Neurobiology of an Insect Brain. Oxford University Press. Oxford University Press

Cadwell, C. R., Palasantza, A., Jiang, X., Berens, P., Deng, Q., Yilmaz, M., et al. (2016). Electrophysiological, transcriptomic and morphologic profiling of single neurons using Patch-seq. Nature Biotechnology 34, 199–203.

Ciabatti, E., González-Rueda, A., Mariotti, L., Morgese, F., and Tripodi, M. (2017). Life-Long Genetic and Functional Access to Neural Circuits Using Self-Inactivating Rabies Virus. Cell 170, 382–392.e14.

Cossell, L., Iacaruso, M. F., Muir, D. R., Houlton, R., Sader, E. N., Ko, H., et al. (2015). Functional organization of excitatory synaptic strength in primary visual cortex. Nature 518, 399–403.

Crochet, S., Chauvette, S., Boucetta, S., and Timofeev, I. (2005). Modulation of synaptic transmission in neocortex by network activities. Eur. J. Neurosci. 21, 1030–1044.

Csicsvari, J., Hirase, H., Czurko, A., and Buzsaki, G. (1998). Reliability and state dependence of pyramidal cell-interneuron synapses in the hippocampus: an ensemble approach in the behaving rat. Neuron 21, 179–189.

Daigle, T. L., Madisen, L., Hage, T. A., Valley, M. T., Knoblich, U., Larsen, R. S., et al. (2018). A Suite of Transgenic Driver and Reporter Mouse Lines with Enhanced Brain-Cell-Type Targeting and Functionality. Cell 174, 465–480.e22.

Debanne, D., Boudkkazi, S., Campanac, E., Cudmore, R. H., Giraud, P., Fronzaroli-Molinieres, L., et al. (2008). Paired-recordings from synaptically coupled cortical and hippocampal neurons in acute and cultured brain slices. Nat Protoc 3, 1559–1568.

Deuchars, J., and Thomson, A. M. (1995). Innervation of burst firing spiny interneurons by pyramidal cells in deep layers of rat somatomotor cortex: paired intracellular recordings with biocytin filling. Neuroscience 69, 739–755.

Edwards, F. A., Konnerth, A., Sakmann, B., and Takahashi, T. (1989). A thin slice preparation for patch clamp recordings from neurones of the mammalian central nervous system. Pfl□gers Archiv European Journal of Physiology 414, 600–612.

English, D. F., McKenzie, S., Evans, T., Kim, K., Yoon, E., and Buzsáki, G. (2017). Pyramidal Cell-Interneuron Circuit Architecture and Dynamics in Hippocampal Networks. Neuron 96, 505–520.e7.

Feldmeyer, D., and Radnikow, G. (2016). “Paired Recordings from Synaptically Coupled Neurones in Acute Neocortical Slices,” in Advanced Patch-Clamp Analysis for Neuroscientists Neuromethods. (New York, NY: Springer New York), 171–191.

Feldmeyer, D., Lübke, J., and Sakmann, B. (2006). Efficacy and connectivity of intracolumnar pairs of layer 2/3 pyramidal cells in the barrel cortex of juvenile rats. J. Physiol. (Lond.) 575, 583–602.

Fernández, A., Radmilovich, M., Russo, R. E., Hounsgaard, J., and Trujillo-Cenóz, O. (1996). Monosynaptic connections between primary afferents and giant neurons in the turtle spinal dorsal horn. Exp Brain Res 108, 347–356.

Ferrarese, L., Jouhanneau, J.-S., Remme, M. W. H., Kremkow, J., Katona, G., Rózsa, B., et al. (2018). Dendrite-Specific Amplification of Weak Synaptic Input during Network Activity In Vivo. Cell Rep 24, 3455–3465.e5.

Fujisawa, S., Amarasingham, A., Harrison, M. T., and Buzsáki, G. (2008). Behavior-dependent short-term assembly dynamics in the medial prefrontal cortex. Nat. Neurosci. 11, 823–833.

Fuzik, J., Zeisel, A., Mate, Z., Calvigioni, D., Yanagawa, Y., Szabó, G., et al. (2016). Integration of electrophysiological recordings with single-cell RNA-seq data identifies neuronal subtypes. Nature Biotechnology 34, 175–183.

Geiger, J. R., Lübke, J., Roth, A., Frotscher, M., and Jonas, P. (1997). Submillisecond AMPA receptor-mediated signaling at a principal neuron-interneuron synapse. Neuron 18, 1009–1023.

Gentet, L. J., Avermann, M., Matyas, F., Staiger, J. F., and Petersen, C. C. H. (2010). Membrane potential dynamics of GABAergic neurons in the barrel cortex of behaving mice. Neuron 65, 422–435.

Gerfen, C. R., Paletzki, R., and Heintz, N. (2013). GENSAT BAC cre-recombinase driver lines to study the functional organization of cerebral cortical and basal ganglia circuits. Neuron 80, 1368–1383.

Goebbels, S., Bormuth, I., Bode, U., Hermanson, O., Schwab, M. H., and Nave, K.-A. (2006). Genetic targeting of principal neurons in neocortex and hippocampus of NEX-Cre mice. genesis 44, 611–621.

Guzman, S. J., Schlögl, A., Frotscher, M., and Jonas, P. (2016). Synaptic mechanisms of pattern completion in the hippocampal CA3 network. Science 353, 1117–1123.

Hamada, H., Takaori, M., Kimura, K., Fukui, A., and Fujita, Y. (1993). Changes in circulating blood volume following isoflurane or sevoflurane anesthesia. J Anesth 7, 316–324.

Harris, J. A., Hirokawa, K. E., Sorensen, S. A., Gu, H., Mills, M., Ng, L. L., et al. (2014). Anatomical characterization of Cre driver mice for neural circuit mapping and manipulation. Front. Neural Circuits 8, 76.

Hippenmeyer, S., Vrieseling, E., Sigrist, M., Portmann, T., Laengle, C., Ladle, D. R., et al. (2005). A developmental switch in the response of DRG neurons to ETS transcription factor signaling. PLoS Biol. 3, e159.

Holmgren, C., Harkany, T., Svennenfors, B., and Zilberter, Y. (2003). Pyramidal cell communication within local networks in layer 2/3 of rat neocortex. J. Physiol. (Lond.) 551, 139–153.

Jiang, X., Shen, S., Cadwell, C. R., Berens, P., Sinz, F., Ecker, A. S., et al. (2015). Principles of connectivity among morphologically defined cell types in adult neocortex. Science 350, aac9462–aac9462.

Jouhanneau, J.-S., Kremkow, J., and Poulet, J. F. A. (2018). Single synaptic inputs drive high-precision action potentials in parvalbumin expressing GABA-ergic cortical neurons in vivo. Nat Commun 9, 1540.

Jouhanneau, J.-S., Kremkow, J., Dorrn, A. L., and Poulet, J. F. A. (2015). In Vivo Monosynaptic Excitatory Transmission between Layer 2 Cortical Pyramidal Neurons. Cell Rep 13, 2098–2106.

Kitamura, K., Judkewitz, B., Kano, M., Denk, W., and Häusser, M. (2008). Targeted patch-clamp recordings and single-cell electroporation of unlabeled neurons in vivo. Nat. Methods 5, 61–67.

Ko, H., Hofer, S. B., Pichler, B., Buchanan, K. A., Sjöström, P. J., and Mrsic-Flogel, T. D. (2011). Functional specificity of local synaptic connections in neocortical networks. Nature 473, 87–91.

Kolb, I., Stoy, W. A., Rousseau, E. B., Moody, O. A., Jenkins, A., and Forest, C. R. (2016). Cleaning patch-clamp pipettes for immediate reuse. Sci Rep 6, 35001.

Lalanne, T., Abrahamsson, T., and Sjöström, P. J. (2016). Using Multiple Whole-Cell Recordings to Study Spike-Timing-Dependent Plasticity in Acute Neocortical Slices. Cold Spring Harb Protoc 2016, pdb.prot091306.

Lee, A. K., and Brecht, M. (2018). Elucidating Neuronal Mechanisms Using Intracellular Recordings during Behavior. Trends in Neurosciences 41, 385–403.

Lefort, S., Tomm, C., Sarria, J. C. F., and Petersen, C. C. H. (2009). The Excitatory Neuronal Networkof the C2 Barrel Columnin Mouse Primary Somatosensory Cortex. Neuron 61, 301–316.

London, M., Roth, A., Beeren, L., Häusser, M., and Latham, P. E. (2010). Sensitivity to perturbations in vivo implies high noise and suggests rate coding in cortex. Nature 466, 123–127.

Madisen, L., Zwingman, T. A., Sunkin, S. M., Oh, S. W., Zariwala, H. A., Gu, H., et al. (2010). A robust and high-throughput Cre reporting and characterization system for the whole mouse brain. Nat. Neurosci. 13, 133–140.

Margrie, T., Brecht, M., and Sakmann, B. (2002). In vivo, low-resistance, whole-cell recordings from neurons in the anaesthetized and awake mammalian brain. Pfl□gers Archiv European Journal of Physiology 444, 491–498.

Markram, H., Lübke, J., Frotscher, M., and Sakmann, B. (1997a). Regulation of synaptic efficacy by coincidence of postsynaptic APs and EPSPs. Science 275, 213–215.

Markram, H., Lübke, J., Frotscher, M., Roth, A., and Sakmann, B. (1997b). Physiology and anatomy of synaptic connections between thick tufted pyramidal neurones in the developing rat neocortex. J. Physiol. (Lond.) 500 (Pt 2), 409–440.

Mason, A., Nicoll, A., and Stratford, K. (1991). Synaptic transmission between individual pyramidal neurons of the rat visual cortex in vitro. J. Neurosci. 11, 72–84.

Matsumura, M., Chen, D., Sawaguchi, T., Kubota, K., and Fetz, E. E. (1996). Synaptic interactions between primate precentral cortex neurons revealed by spike-triggered averaging of intracellular membrane potentials in vivo. J. Neurosci. 16, 7757–7767.

Muñoz, W., Tremblay, R., Levenstein, D., and Rudy, B. (2017). Layer-specific modulation of neocortical dendritic inhibition during active wakefulness. Science 355, 954–959.

Packer, A. M., Russell, L. E., Dalgleish, H. W. P., and Häusser, M. (2015). Simultaneous all-optical manipulation and recording of neural circuit activity with cellular resolution in vivo. Nat. Methods 12, 140–146.

Pala, A., and Petersen, C. C. (2018). State-dependent cell-type-specific membrane potential dynamics and unitary synaptic inputs in awake mice. eLife 7, 350.

Palmer, L. M., Shai, A. S., Reeve, J. E., Anderson, H. L., Paulsen, O., and Larkum, M. E. (2014). NMDA spikes enhance action potential generation during sensory input. Nat. Neurosci. 17, 383–390.

Papadopoulos, I. N., Jouhanneau, J.-S., Poulet, J. F. A., and Judkewitz, B. (2017). Scattering compensation by focus scanning holographic aberration probing (F-SHARP). Nature Photonics 2016 11:2 11, 116–123.

Parker, D. (2003). Variable properties in a single class of excitatory spinal synapse. J. Neurosci. 23, 3154–3163.

Parker, D. (2010). Neuronal network analyses: premises, promises and uncertainties. Philosophical Transactions of the Royal Society B: Biological Sciences 365, 2315–2328.

Peng, Y., Barreda Tomás, F. J., Klisch, C., Vida, I., and Geiger, J. R. P. (2017). Layer-Specific Organization of Local Excitatory and Inhibitory Synaptic Connectivity in the Rat Presubiculum. Cereb. Cortex 27, 2435–2452.

Perin, R., Berger, T. K., and Markram, H. (2011). A synaptic organizing principle for cortical neuronal groups. Proc. Natl. Acad. Sci. U.S.A. 108, 5419–5424.

Petersen, C. C. H. (2017). Whole-Cell Recording of Neuronal Membrane Potential during Behavior. Neuron 95, 1266–1281.

Poulet, J. F. A., and Hedwig, B. (2006). The cellular basis of a corollary discharge. Science 311, 518–522.

Poulet, J. F. A., and Petersen, C. C. H. (2008). Internal brain state regulates membrane potential synchrony in barrel cortex of behaving mice. Nature 454, 881–885.

Reardon, T. R., Murray, A. J., Turi, G. F., Wirblich, C., Croce, K. R., Schnell, M. J., Jessel, T. M., and Losonczy, A. (2016). Rabies Virus CVS-N2c(ΔG) Strain Enhances Retrograde Synaptic Transfer and Neuronal Viability. Neuron. 89, 711–724.

Reid, R. C., and Alonso, J. M. (1995). Specificity of monosynaptic connections from thalamus to visual cortex. Nature 378, 281–284.

Rickgauer, J. P., Deisseroth, K., and Tank, D. W. (2014). Simultaneous cellular-resolution optical perturbation and imaging of place cell firing fields. Nat. Neurosci. 17, 1816–1824.

Roberts, A., Li, W.-C., and Soffe, S. R. (2010). How neurons generate behavior in a hatchling amphibian tadpole: an outline. Front Behav Neurosci 4, 16.

Sofroniew, N. J., Flickinger, D., King, J., and Svoboda, K. (2016). A large field of view two-photon mesoscope with subcellular resolution for in vivo imaging. eLife 5, 413.

Steriade, M., Nuñez, A., and Amzica, F. (1993). A Novel Slow (<1 Hz) Oscillation of Neocortical Neurons in viva: Depolarizing and Hyperpolarizing Components. Journal of Neuroscience Methods 13, 3252–3265.

Stirman, J. N., Smith, I. T., Kudenov, M. W., and Smith, S. L. (2016). Wide field-of-view, multi-region, two-photon imaging of neuronal activity in the mammalian brain. Nature Biotechnology 34, 857–862.

Suk, H.-J., van Welie, I., Kodandaramaiah, S. B., Allen, B., Forest, C. R., and Boyden, E. S. (2017). Closed-Loop Real-Time Imaging Enables Fully Automated Cell-Targeted Patch-Clamp Neural Recording In Vivo. Neuron 96, 244–245.

Swadlow, H. A., and Gusev, A. G. (2002). Receptive-field construction in cortical inhibitory interneurons. Nat. Neurosci. 5, 403–404.

Tamamaki, N., Yanagawa, Y., Tomioka, R., Miyazaki, J.-I., Obata, K., and Kaneko, T. (2003). Green fluorescent protein expression and colocalization with calretinin, parvalbumin, and somatostatin in the GAD67-GFP knock-in mouse. J. Comp. Neurol. 467, 60–79.

Taniguchi, H., He, M., Wu, P., Kim, S., Paik, R., Sugino, K., et al. (2011). NeuroResource. Neuron 71, 995–1013.

Urban-Ciecko, J., Jouhanneau, J.-S., Myal, S. E., Poulet, J. F. A., and Barth, A. L. (2018). Precisely Timed Nicotinic Activation Drives SST Inhibition in Neocortical Circuits. Neuron 97, 611–625.e5.

Vélez-Fort, M., Bracey, E. F., Keshavarzi, S., Rousseau, C. V., Cossell, L., Lenzi, S. C., et al. (2018). A Circuit for Integration of Head- and Visual-Motion Signals in Layer 6 of Mouse Primary Visual Cortex. Neuron 98, 179–191.e6.

Wang, G., Wyskiel, D. R., Yang, W., Wang, Y., Milbern, L. C., Lalanne, T., et al. (2015). An optogenetics- and imaging-assisted simultaneous multiple patch-clamp recording system for decoding complex neural circuits. Nat Protoc 10, 397–412.

Weiler, S., Bauer, J., Hübener, M., Bonhoeffer, T., Rose, T., and Scheuss, V. (2018). High-yield in vitro recordings from neurons functionally characterized in vivo. Nat Protoc 13, 1275–1293.

Wertz, A., Trenholm, S., Yonehara, K., Hillier, D., Raics, Z., Leinweber, M., et al. (2015). PRESYNAPTIC NETWORKS. Single-cell-initiated monosynaptic tracing reveals layer-specific cortical network modules. Science 349, 70–74.

Wickersham, I. R., Lyon, D. C., Barnard, R. J. O., Mori, T., Finke, S., Conzelmann, K.-K., et al. (2007). Monosynaptic restriction of transsynaptic tracing from single, genetically targeted neurons. Neuron 53, 639–647.

Wozny, C., and Williams, S. R. (2011). Specificity of synaptic connectivity between layer 1 inhibitory interneurons and layer 2/3 pyramidal neurons in the rat neocortex. Cereb. Cortex 21, 1818–1826.

Yassin, L., Benedetti, B. L., Jouhanneau, J.-S., Wen, J. A., Poulet, J. F. A., and Barth, A. L. (2010). An Embedded Subnetwork of Highly Active Neurons in the Neocortex. Neuron 68, 1043–1050.

Yu, J., and Ferster, D. (2013). Functional coupling from simple to complex cells in the visually driven cortical circuit. J. Neurosci. 33, 18855–18866.

Zhao, W.-J., Kremkow, J., and Poulet, J. F. A. (2016). Translaminar Cortical Membrane Potential Synchrony in Behaving Mice. Cell Rep 15, 1–34.

